# Three-dimensional Imaging of Colonial Cyanobacteria with Optical Coherence Tomography

**DOI:** 10.64898/2026.08.27.747059

**Authors:** Yuri Z. Sinzato, Robert Uittenbogaard, Petra M. Visser, Jef Huisman, Maziyar Jalaal

**Affiliations:** Van der Waals-Zeeman Institute, Institute of Physics, University of Amsterdam, Science Park 904, Amsterdam, 1098XH, The Netherlands; Hydro-Key Ltd, Haelen 6081EA, The Netherlands; Department of Freshwater and Marine Ecology, Institute for Biodiversity and Ecosystem Dynamics, University of Amsterdam, Science Park 904, Amsterdam, 1098XH, The Netherlands; Department of Applied Mathematics and Theoretical Physics, University of Cambridge, Wilberforce Road, Cambridge CB3 0WA, United Kingdom

**Keywords:** Algal blooms, Morphology, Visualization, Cyanobacterial colonies, *Microcystis*

## Abstract

The morphology of cyanobacterial colonies plays a key role in harmful cyanobacterial blooms, with implications for their vertical migration, resistance against grazing, and light availability. In this study, we introduce the use of Optical Coherence Tomography (OCT) to investigate the three-dimensional morphology of cyanobacterial colonies. The technique enables non-invasive 3D imaging of colonies up to several millimeters in size, providing access to detailed mesoscale morphological features. Gas vesicles inside cells were shown to strongly improve image quality. We describe the sample preparation and image acquisition protocol, as well as an image processing pipeline that extracts mesoscale morphological features and provides a volumetric visualization of colonies. The method was tested for representative colonies of different cyanobacterial species while a dataset of volumetric images and measured mesoscale features was acquired for natural colonies of *Microcystis*. We demonstrate the utility of 3D imaging by quantifying the effects of irregular colony morphologies on their flotation velocity and the light availability within colonies. We anticipate OCT to become a key imaging technique to monitor populations of cyanobacterial colonies and investigate colony formation, with potential extensions to other colonial and aggregated organisms in freshwater and marine environments.

## I. INTRODUCTION

Colony formation is a key trait of many bloom-forming cyanobacteria (Reynolds, 2006; Huisman et al., 2018; Xiao et al., 2018). These colonies exhibit diverse architectures: *Microcystis* typically forms spherical and branched clusters ranging in size from a few cells to millions of cells, filamentous genera such as *Aphanizomenon* often form raft-like structures in which the filaments are aligned in parallel, whereas *Dolichospermum* and *Gloeotrichia* form entangled filamentous aggregates of varied geometrical shapes. Cyanobacterial colonies can be remarkably resistant to hydrodynamic stress (Sinzato et al., 2026a), and colonial morphology strongly influences both the physiological functioning and population dynamics of cyanobacteria (Fig. 1). For example, colony size and shape influence the flotation velocity of colonies (Nakamura et al., 1993; Li et al., 2016; Sinzato et al., 2026b), which in turn affects their vertical distribution and light access in the water column (Ibelings et al., 1991; Visser et al., 1997; Huisman et al., 2004). Colonial morphology also modulates light penetration into the interior of the colonies through self-shading (Feng et al., 2019, 2024), with potential consequences for photosynthesis and buoyancy regulation (Xu et al., 2023). In addition, the shape of cyanobacterial colonies can affect protection against grazing by zooplankton (Yang et al., 2006; Cerbin et al., 2013; Lürling, 2021). Among colonial cyanobacteria, *Microcystis* in particular has been studied extensively due to its global prevalence (Harke et al., 2016) and its ability to produce dense and often toxic blooms (Huisman et al., 2018). The morphology of *Microcystis* colonies is also important from a taxonomic perspective. For example, structural features such as branching and elongation are commonly used to classify different morphospecies of *Microcystis* (Komárek & Komárková, 2002) and to monitor seasonal variation in morphospecies composition (Feng et al., 2020), although the marked morphological plasticity of this genus complicates such classifications (Xiao et al., 2018).

**Fig. 1:**
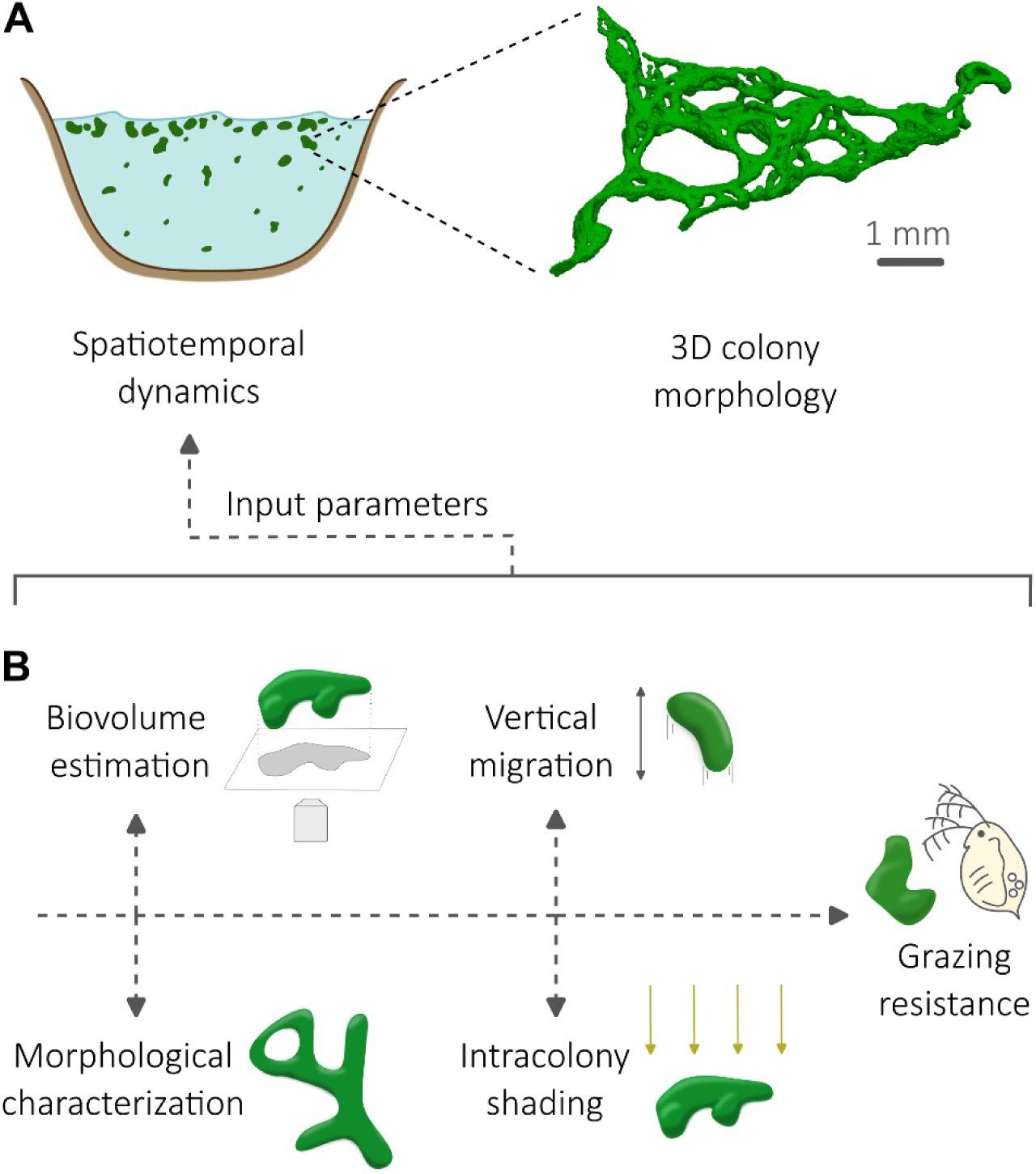
Ecological relevance of the three-dimensional morphological features of cyanobacterial colonies. (A) Complex three-dimensional morphology of a large colony of *Microcystis aeruginosa* imaged with OCT and depicted by a volumetric visualization. Such complex geometries influence the spatiotemporal dynamics of cyanobacterial colonies in a water column. (B) Schematics of a set of traits associated with different morphological features.

Given the central role of colonial morphology in the physiology and ecology of bloom-forming cyanobacteria, there is a clear need for imaging techniques that can resolve their full three-dimensional structure while requiring minimal sample preparation. However, current imaging approaches remain insufficient to fully characterize the complex 3D architecture of cyanobacterial colonies. The 2D images obtained by wide field microscopy provide useful information on cell size, intercellular spacing, and extracellular polymeric substances (EPS), but depth information is limited to colony projections along the imaging axis (Sampognaro et al., 2020; Bormans et al., 2023). Current methods for estimating colony biovolume rely on assumptions of colony shape or empirical relations between cell number and colony size (Li et al., 2014; Alcántara et al., 2018; T-Krasznai et al., 2022). Confocal fluorescence microscopy (Mougin et al., 2025) can measure the axial position of cells along the imaging axis, but the imaging depth is limited to a few cell layers and its application to colonies treated with fixatives (e.g., Lugol’s iodine) is often hampered by the loss of chlorophyll fluorescence. More broadly, there is a need for dedicated 3D imaging approaches that are tailored to study colonial morphologies across a wide range of cyanobacterial genera. Optical coherence tomography (OCT) offers a great potential to image the mesoscale features (i.e., from a few cells in length up to the colony size) of cyanobacterial colonies. OCT is a non-invasive and high-resolution imaging technique invented in the early 1990s (Huang et al., 1991) that uses interferometry to acquires 2D and 3D images of biological tissues and other scattering media (Fercher et al., 2003; Podoleanu, 2012; Bouma et al., 2022). OCT has been successfully applied to investigate the structure and dynamical behavior of bacterial biofilms (Haisch & Niessner, 2007; Wagner & Horn, 2017; Picioreanu et al., 2018), including marine cyanobacterial biofilms (Romeu et al., 2019; Faria et al., 2021). However, those studies were limited to substrate-attached benthic cyanobacteria.

The operating range of the OCT system, defined here as the range from lateral resolution up to the field of view (Fig. 2A), makes it ideal for imaging mesoscale features of bacterial colonies, which can reach dimensions of a few cells up to large colonies of several millimeters in size. This operating range is complementary to other 3D imaging techniques commonly used in cellular and microbial research (Fig. 2A), such as confocal fluorescence microscopy (Mougin et al., 2025) for measurements of the size and shape of individual cells and intercellular arrangements, and transmission electron microscopy (Reynolds et al., 1981; Gaëtan et al., 2023) for visualizing intracellular structures such as gas vesicles in cyanobacteria or specific organelles in eukaryotic organisms.

**Fig. 2:**
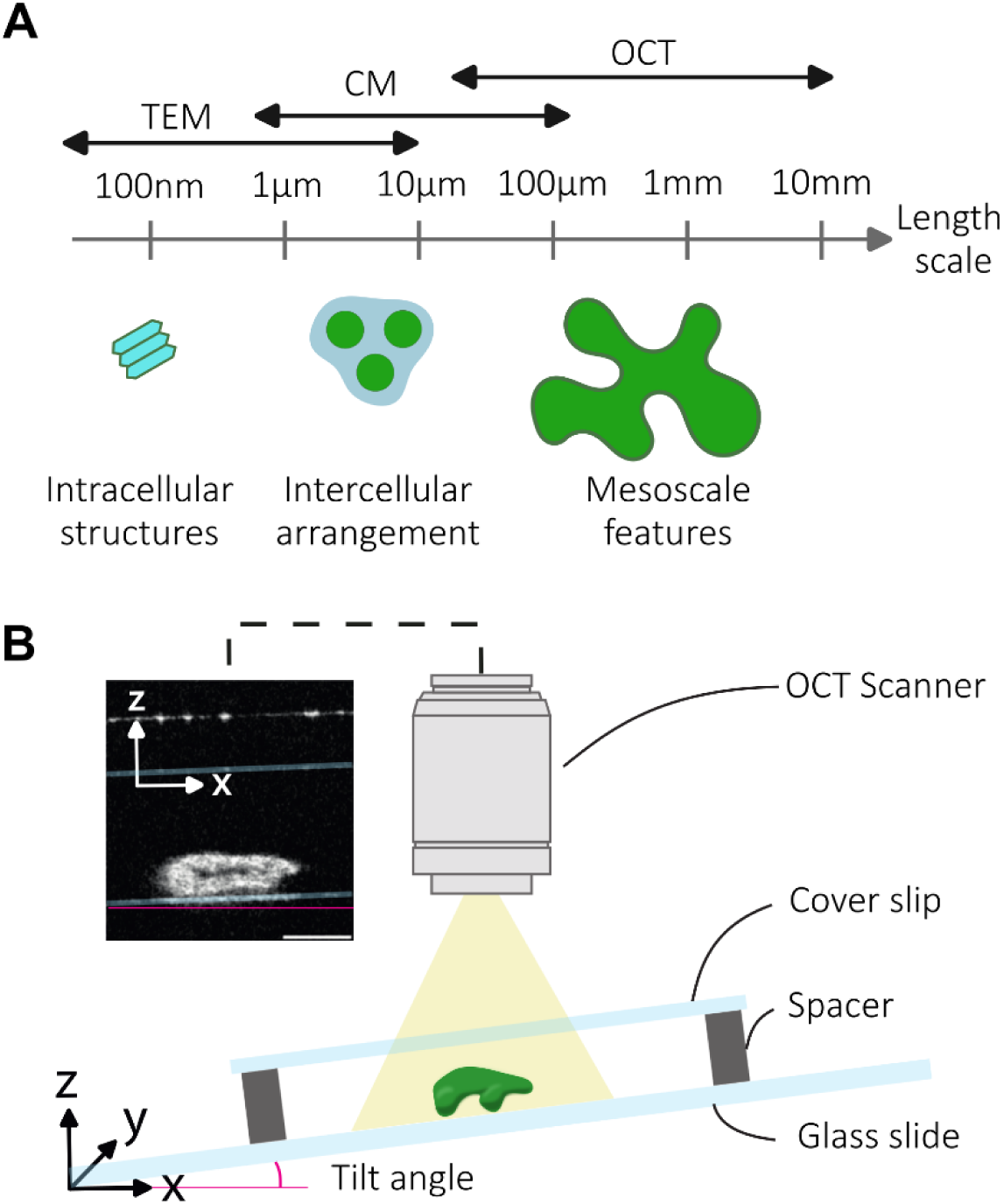
Schematic of the setup for three-dimensional imaging of cyanobacterial colonies with optical coherence tomography (OCT). (A) Operating range for different three-dimensional imaging techniques applied to cyanobacterial colonies. The double arrows indicate the typical length scales from the smallest lateral resolution to the largest field of view covered by each technique. Optical coherence tomography depicts mesoscale features, while confocal fluorescence microscopy (CM) depicts intercellular arrangement and cellular shape and size, and transmission electron microscopy (TEM) depicts intracellular structures such as gas vesicles. (B) A suspension of colonies is enclosed in an imaging chamber consisting of two glass surfaces separated by a spacer. The chamber is placed at a tilt angle with respect to the imaging axis of the OCT scanner in order to reduce reflection from the glass. Inset is an example of OCT vertical slice (XZ) from a colony of *Microcystis ichthyoblabe* sampled from Lake Braassemermeer in 2025. Scale bar indicates 200 µm.

Extending OCT to buoyant cyanobacterial colonies is especially compelling because their cells contain gas vesicles, which act as natural optical contrast agents and promote strong light scattering (Dubelaar et al., 1987; Walsby, 1991; Pfeifer, 2012). In fact, gas vesicles from cyanobacteria have been isolated for use as contrast agents for ultrasound and magnetic resonance imaging (Lakshmanan et al., 2017). In a recent study, we have employed OCT imaging to correct for projection-bias in the measurement of the flotation velocity of *Microcystis* spp. colonies (Sinzato et al., 2026b). Nonetheless, a standardized protocol for OCT imaging of suspended colonies of freshwater cyanobacteria is currently lacking. Furthermore, automation of image processing can increase the throughput of colonies and reduce subjective input from the operator (Hou et al., 2019).

In this study, we have developed a protocol for 3D image acquisition of cyanobacterial colonies using OCT. We have also developed an automated image processing pipeline for the measurement of mesoscale features and volumetric visualization of colonies. We assessed the impact of gas vesicles in the OCT signal intensity and compared the results to confocal fluorescence microscopy. The protocol was tested for field-collected colonies of different cyanobacterial taxa. An extensive dataset of OCT scans and measurements of mesoscale morphological features was acquired for colonies of *Microcystis* spp. Finally, we used the volumetric images of *Microcystis* colonies to estimate their flotation velocity and intracolony light attenuation and compare the results obtained by OCT for these 3D colonial shapes with simplified approaches assuming idealized spherical colonies.

## II. METHODS AND PROCEDURES

### A. Sample collection and preparation

Cyanobacterial colonies were collected from several lakes in the Netherlands in September 2024 and between June and October 2025 (see Supplementary Table S1 for location and date of each sample). Plankton samples were concentrated with a 100 µm net and transported in cooled vials to the laboratory within 2 hours of collection. The vials were centrifuged (100 g, 15 min) such that sediment and non-buoyant plankton were collected in the pellet. The supernatant containing a suspension of buoyant cyanobacterial colonies was separated for imaging. The suspension of buoyant colonies was stored at 4 °C in the dark for 1 to 5 days until imaging. Immediately before OCT image acquisition, the suspension of colonies was gently injected in an imaging chamber (Fig. 2B). The chamber consisted of two glass surfaces (a microscope slide of 75 x 25 x 1 mm and a cover slip of 20 x 20 x 0.13-0.16 mm) separated by a spacer (one or multiple layers of hand-cut double sided tape) with sufficient height to enclose the colonies without squeezing (chambers varied from 0.5 to 2 mm in spacer height). Care was taken to avoid the presence of bubbles inside the imaging chamber. Each chamber contained from a few up to 100 colonies; the number of colonies was limited to avoid overlap of neighboring colonies during imaging.

A thick *Dolichospermum* sp. colony (height > 300 µm) was used to assess the effect of gas vesicles on the OCT image, as the large thickness allowed quantification of the signal along the colony height. The colony with intact gas vesicles was first imaged. Subsequently, the chamber containing the colony was inserted in a pressure vessel and 1 MPa of nitrogen gas was applied for a few seconds, after which the gas vesicles inside cells collapsed (Walsby et al., 1992). The colony with collapsed gas vesicles was again imaged with OCT.

Representative colonies of different cyanobacterial taxa were selected to assess the performance of OCT. Each of the selected colonies was also imaged with confocal fluorescence microscopy and bright field microscopy. Filamentous cyanobacteria were identified to genus level from the bright field microscopy images and morphospecies of *Microcystis* were identified following the criteria proposed by Komárek & Komárková (2002). A large number of buoyant *Microcystis* spp. colonies was collected from Lake ’t Joppe in September 2024 (sample J-09-24) and August 2025 (sample J-08-25; Supplementary Table S1). Subsamples of these colony suspensions were injected into imaging chambers and all colonies larger than 100 µm in the chambers were scanned with OCT (N = 65 colonies in 2024 and N = 148 colonies in 2025). We were unable to track the morphospecies identity of individual colonies within the suspension, as the individual colonies scanned with OCT could not be located again during microscopy. Therefore, another subsample of the colony suspensions was injected in a cylindrical counting chamber and morphospecies composition was identified by bright field microscopy.

### B. Optical coherence tomography

#### B.1. Image acquisition

The working principles of OCT and the different design configurations have been described by several reviews (Fercher et al., 2003; Podoleanu, 2012; Bouma et al., 2022). In this study, we have used a spectral-domain OCT imaging system (Telesto 221C1/M equipped with a LK3 scan lens, Thorlabs GmbH, Germany). The sample was illuminated with a low-coherence light in the near-infrared region (1300 nm). The light is back-scattered or back-reflected by the sample into the scanner. The image is reconstructed from the phase delay between the sample light path and a reference light path. A one-dimensional scan along the imaging axis (here defined as *z*) is commonly named A-scan (Wagner & Horn, 2017). A sequence of A-scans along a transverse direction (*x*) forms a cross-section of the sample, named B-scan. Proceeding with a sequence of B-scans along the third orthogonal axis (*y*) generates a 3D scan of the sample. A typical 3D scan of 400 x 400 x 400 pixels had a total acquisition time of ∼ 10 s. Using common conventions for OCT measurements, pixel values in the OCT scan are reported as the signal-to-noise ratio (SNR), *i.e.*, the ratio between signal intensity (S) and the background intensity (B) measured on a logarithmic scale and expressed as decibels, SNR = 10 log_10_ (S/B).

The axial resolution of our OCT system was 4.2 µm and the lateral resolution was 13 µm. The maximum field of view was 10 x 10 mm, while the maximum imaging depth was at least 2.0 mm, which is larger than all cyanobacterial colonies present in our samples. To acquire the scans, the imaging chamber containing the sample was placed below the OCT scanner with a tilt angle of 10° between the imaging axis and the normal vector of the flat glass surfaces (Fig. 2B). The chamber tilt reduces light reflection from the glass surfaces into the scanner. The colonies were kept at a distance of at least 10 % of the image height from the top (maximum z) and bottom (minimum z) edges of the image, as the peripheral regions of the OCT image have a lower signal-to-noise ratio.

#### B.2. Image processing

The 3D scans of cyanobacterial colonies underwent several image processing steps to remove artifacts, detect colony regions, correct for chamber tilt and measure morphological features. We developed an automated pipeline to perform image processing (Fig. 3). The script for the pipeline was written in jupyter notebook and was based on the library scikit-image (Van der Walt et al., 2014). The script is accessible at https://github.com/FluidLab/CyanOCT together with detailed instructions. A raw scan consisted of a stack of vertical slices (*XZ*) along the y-axis (Fig. 3A). First, a Gaussian blur filter was applied to reduce noise (Fig. 3B). Second, a straight line Hough transform was applied to detect the glass surfaces (Fig. 3C) and measure the tilt angle. The region of interest (ROI) was selected as the region between the two glass surfaces and a binary mask was created, where the pixels inside the ROI are set to 1, while the pixels outside the ROI are set to 0. The binary mask (Fig. 3C) and the denoised image (Fig. 3B) were multiplied (element-wise) to create a masked image (Fig. 3D) without the glass surfaces. Colony pixels were separated from background pixels with an Otsu-thresholding method. Binary closing and opening are sequentially applied to remove small holes and bright spots smaller than lateral resolution. The colony regions in the resulting thresholded image were segmented with connected-component labeling (Fig. 3E). The <u>labeled</u> image was rotated to correct for the tilt angle, such that the glass surface was normal to the z-axis (Fig. 3F). The final output was a 3D binary image of each colony, in which pixels that belong to the colony have value 1 and pixels outside of the colony have value 0. Volumetric visualization of the colonies was rendered from the binary image with tomviz (Fig. 3G), an open-source application for 3D tomographic data (Levin et al., 2018).

**Fig. 3:**
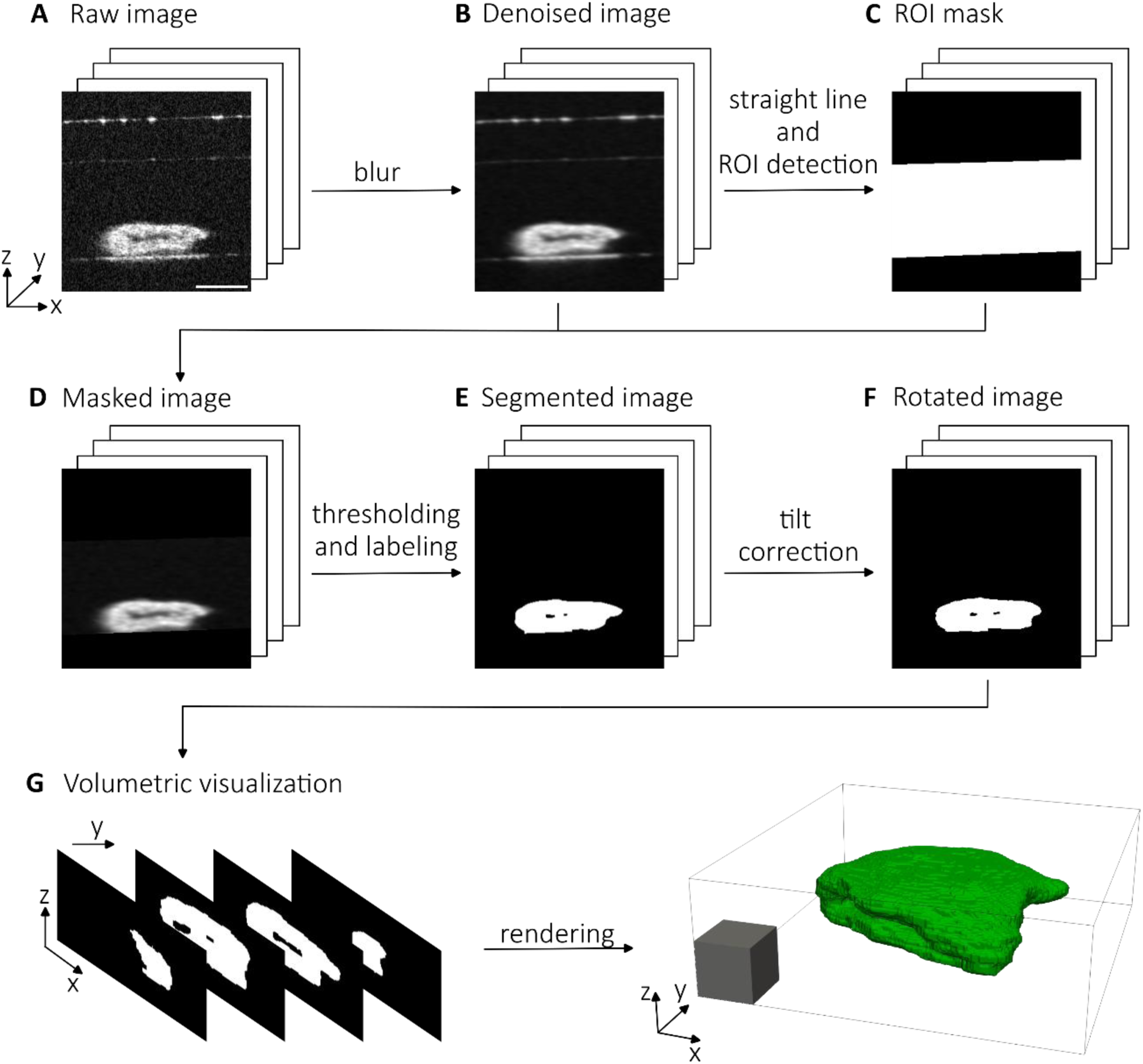
Image processing pipeline to convert raw three-dimensional scans of optical coherence tomography into labeled images. (A) Raw images consist of a stack of vertical slices (XZ) along the Y-axis. Scale bar indicates 200 µm. (B) Noise is reduced using a Gaussian blur filter. (C) The region of interest (ROI), consisting of the space between the two glass surfaces, is detected using a straight line Hough transform. (D) The regions outside the ROI are removed using a masking operation. (E) The colonies are separated from the background with an Otsu-threshold and each colony is segmented using connected-component labelling. (F) The tilt in the image with respect to the imaging axis is corrected with image rotation. (G) The binary vertical slices are rendered into a volumetric visualization. Scale cube has 100 µm of side length. This example is based on a colony of *Microcystis ichthyoblabe* sampled from Lake Braassemermeer in 2025.

#### B.3. Measurement of mesoscale morphological features

Mesoscale morphological features were measured from the OCT scans of *Microcystis* spp. colonies collected from Lake ‘t Joppe in September 2024 (sample J-09-24) and August 2025 (sample J-08-25). The morphological features were measured from the 3D binary images of the colonies using an image processing script based on the python library scikit-image (Van der Walt et al., 2014). The size of *Microcystis* colonies can be quantified by different definitions of colony diameter. From the total volume of the colony, *V_c_*, we define the equivalent spherical diameter *D_es_* as *D_es_* = (6*V_c_/π*)^1*/*3^. The projection of the colony along the *z*-axis has an area *A_z_*, which can be used to define the equivalent circular diameter, *D_ec_* = (4*A_z_/π*)^1*/*2^. Another common measure of colony size is the Feret diameter, *D_f_*, defined as the maximum linear distance between any two points of the colony. For spherical colonies without holes, all three definitions of colony diameter are equivalent, but for nonspherical colonies the different definitions yield different values.

In addition to colony size, the colony structure was characterized by the number of voids (i.e., fully enclosed holes), tunnels (i.e., open holes), and branches per colony. Branches were identified with morphological skeletonization (Saha et al., 2016), which is a commonly used image processing technique to quantify the structure of bacterial biofilms (Geisel et al., 2022; Li et al., 2022). Our skeletonization algorithm (Van der Walt et al., 2014) sequentially removed border pixels from the colony until it was reduced to a network of branches and nodes, referred to as the skeleton (example of colony skeleton given in Fig. S1 and Supplementary Movie S1). The number and length of each branch were computed with the python library skan (Nunez-Iglesias et al., 2018). The thickness of each branch was defined as twice the average distance between the pixels in the branch and their nearest border pixels. Colony voids were considered as part of the colony in the skeletonization algorithm. Terminal branches shorter than their thickness were removed and the average branch thickness of a colony was defined as twice the average distance between all skeleton pixels and their nearest border pixels.

### C. Flotation velocity estimate

The flotation velocity *U* of each *Microcystis* colony can be estimated from Stokes’ law (Reynolds 2006):

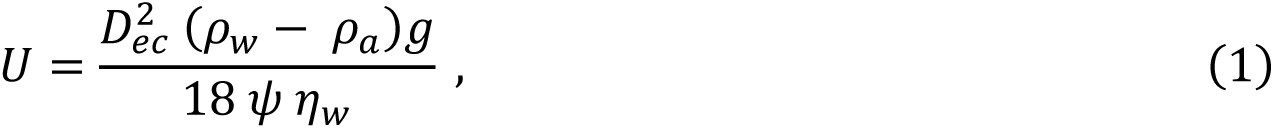

where *g* is the gravitational acceleration, and *ρ_w_* and *η_w_* are the density and dynamic viscosity of water, respectively. The shape factor *ψ* of the colony, which accounts for the effect of colony shape, can be estimated as 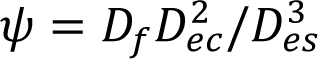 (Sinzato et al., 2026b), where the Feret diameter *D_f_*, the equivalent circular diameter *D_ec_* and the equivalent spherical diameter *D_es_* were measured from the 3D binary image of each colony. The density *ρ_a_* of the colony is a function of the amount of carbohydrate storage, gas vesicles, and EPS within the colony. Since we are here interested in effects of colony morphology rather than colony density, our calculations assumed the same value of *ρ_a_* = 0.988 g*/*mL for all colonies (Sinzato et al., 2026b).

### D. Light attenuation analysis

Irregular colony morphologies can impact the light availability within *Microcystis* colonies. We exemplify this effect with an extension of the intracolony light attenuation model proposed by Feng et al. (2019). Consider a uniform flat layer of *Microcystis* colonies with a thickness *b*. Let *I_o_* be the light intensity incident on the top surface of the layer. Experiments from Feng et al. (2019) demonstrated that the light intensity *I* transmitted through the layer follows Lambert-Beer law:

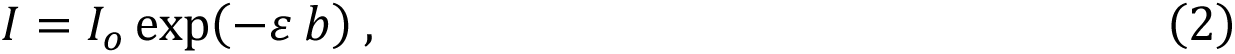

Here, *ε* is the extinction coefficient of the layer, measured at *ε* = 0.045 µm^−1^ for *M. ichthyoblabe* colonies and transmittance above *I/I_o_ >* 0.005 (Feng et al., 2019). We extend this model using 3D binary images of *Microcystis* spp. colonies. Let 1(*r*) denote an indicator function with 1 (***r***) = 1 for ***r*** inside the colony, and 0 for ***r*** outside, where ***r*** = (*x,y,z*) denotes the spatial coordinates. The colony is under a directional incident light intensity *I_o_* oriented along the unit vector ***n***. Applying Lambert-Beer law (Eq. 2), the light intensity *I_n_*(*r*) inside the colony will be

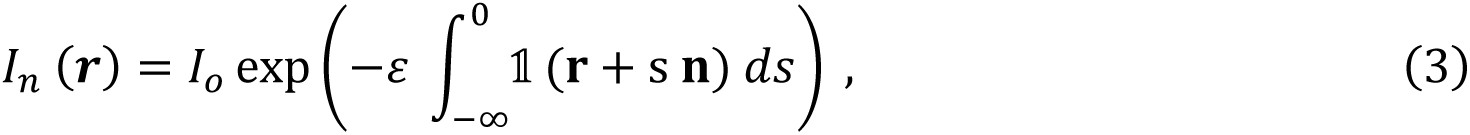

where *s* is an integration variable. Since a colony in suspension may assume any orientation with respect to the light direction, the light intensity in the colony is averaged over all orientations, such that

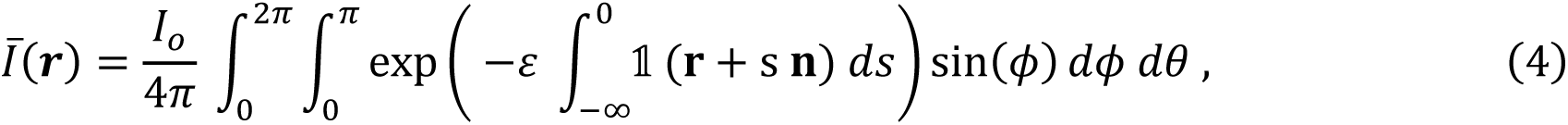

where the orientation unit vector ***n*** = (sin *ϕ* cos *θ ,* sin *ϕ* sin *θ ,* cos *ϕ*) is parametrized by the angles *ϕ* and *θ*. The surface integral in Eq. (4) is computed numerically with midpoint rule integration. An example of the predicted distribution of light intensity inside a *Microcystis* colony is shown in Fig. S2.

### E. Confocal and bright field microscopy

Confocal fluorescence microscopy was used to complement the morphological characterization at the cellular length scale for a subset of cyanobacterial colonies. The imaging chamber containing the colonies was placed in a spinning disk confocal system (Cicero, Visitron Systems GmbH, Germany) and z-stacks were acquired using the fluorescence of chlorophyll *a* at 640 nm excitation. Bright field microscopy images of the cyanobacterial colonies were also acquired using the same imaging system. Maximum intensity projections for the confocal microscopy images were generated with the software ImageJ.

## III. ASSESSMENT & RESULTS

### A. Impact of gas vesicles

As the working principle of OCT is based on back-scattering of light (Fig. 4A) (Fercher et al., 2003), we have compared the OCT signal intensity for a colony of *Dolichospermum* sp. before and after the collapse of gas vesicles, which act as sources of light scattering (Fig. 4B). The signal intensity is reported as the signal-to-noise ratio (*SNR_XY_*) between the average image intensity over a horizontal slice (XY) and the background intensity. The OCT scan of the colony with intact gas vesicles had a maximum *SNR_XY_* of 14 dB. However, after the collapse of gas vesicles, the signal-to-noise ratio dropped over the entire depth of the colony and reached a maximum of 6 dB. In contrast, the fluorescence signal intensity of the same colony measured from the confocal microscopy image displayed an opposite trend (Fig. 4C). The colony with collapsed gas vesicles had a stronger fluorescent intensity, while the intact gas vesicles reduced the signal intensity. However, the effect of gas vesicle collapse on the fluorescence signal was mild (less than 1 dB gain) compared to the strong decrease in signal in the OCT scan. The collapse of gas vesicles was also noticeable from bright field microscopy images of the colony, where the colony with intact gas vesicles was darker than the same colony with collapsed gas vesicles (Fig. 4D). Although the OCT scan with intact gas vesicles yielded a much stronger signal, the OCT scan of the cyanobacterial colony with collapsed gas vesicles still had sufficient intensity to distinguish colony regions from the background along the entire Z axis (Fig. 4E). In contrast, the depth limitation of the confocal microscopy scans was noticeable from their cross-sectional slices, as cells at the top of the colony (thus further from the objective lens of the microscope) displayed a much weaker fluorescence signal than cells at the bottom of the colony (Fig. 4F).

**Fig. 4:**
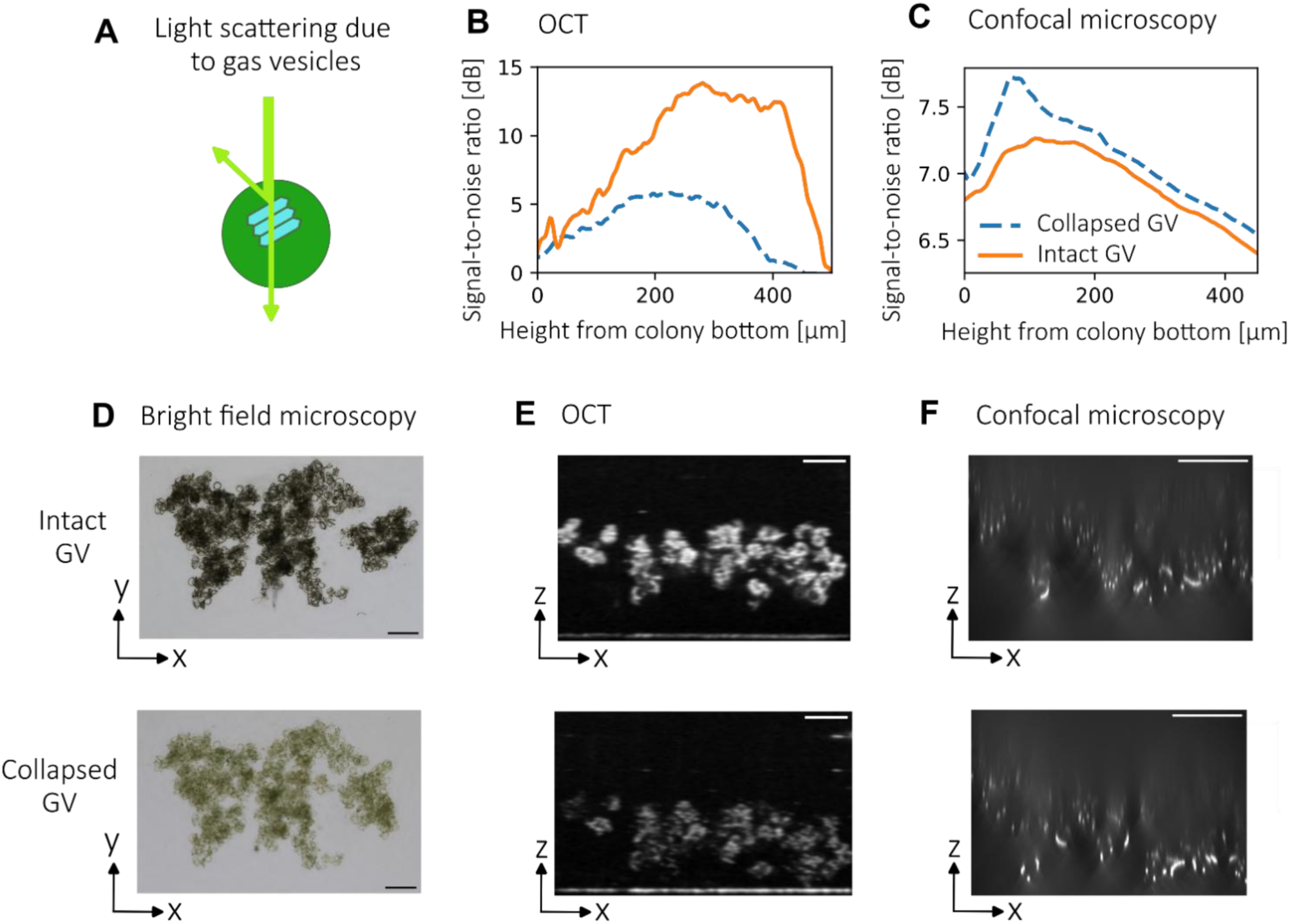
Impact of gas vesicles (GV) on the signal intensity of a *Dolichospermum* sp. colony scanned with optical coherence tomography (OCT) and confocal fluorescence microscopy. (A) Schematics of the light scattering due to the presence of GV inside the cells. (B-C) Signal-to-noise ratio (*SNR_XY_*) between the image intensity averaged over the horizontal XY plane and the background intensity, expressed as decibels (dB), was measured as a function of the height from the colony bottom. The SNR was acquired with (B) OCT and (C) confocal microscopy, where the solid orange line represents the signal with intact GV and the dashed blue line represents the SNR of the same colony after the collapse of GV. (D) The colony with intact GV appeared darker in a bright field microscopy image, while the same colony appeared lighter after the collapse of gas vesicles. (E) Intact GV produced a strong OCT signal intensity along the entire colony, visible from a vertical slice (XZ), while the signal was weaker for collapsed GV. (F) In contrast, intact GV attenuated the image signal in a vertical slice acquired with confocal microscopy, while collapsed GV allowed a higher fraction of the fluorescence signal to be transmitted. Scale bars in panels D-F indicate 200 µm.

### B. Validation for different species of colonial cyanobacteria

Three-dimensional OCT scans were acquired for cyanobacterial colonies of different species (Fig. 5, Supplementary Movie S2.). For all OCT scans, the entire colony could be visualized. Mesoscale features typical of each morphospecies are qualitatively distinguishable in the volumetric visualization of colonies. The OCT scans are complemented with bright field microscopy and confocal microscopy images of the same colonies, where confocal microscopy images are maximum intensity projections of a smaller region of each colony.

**Fig. 5:**
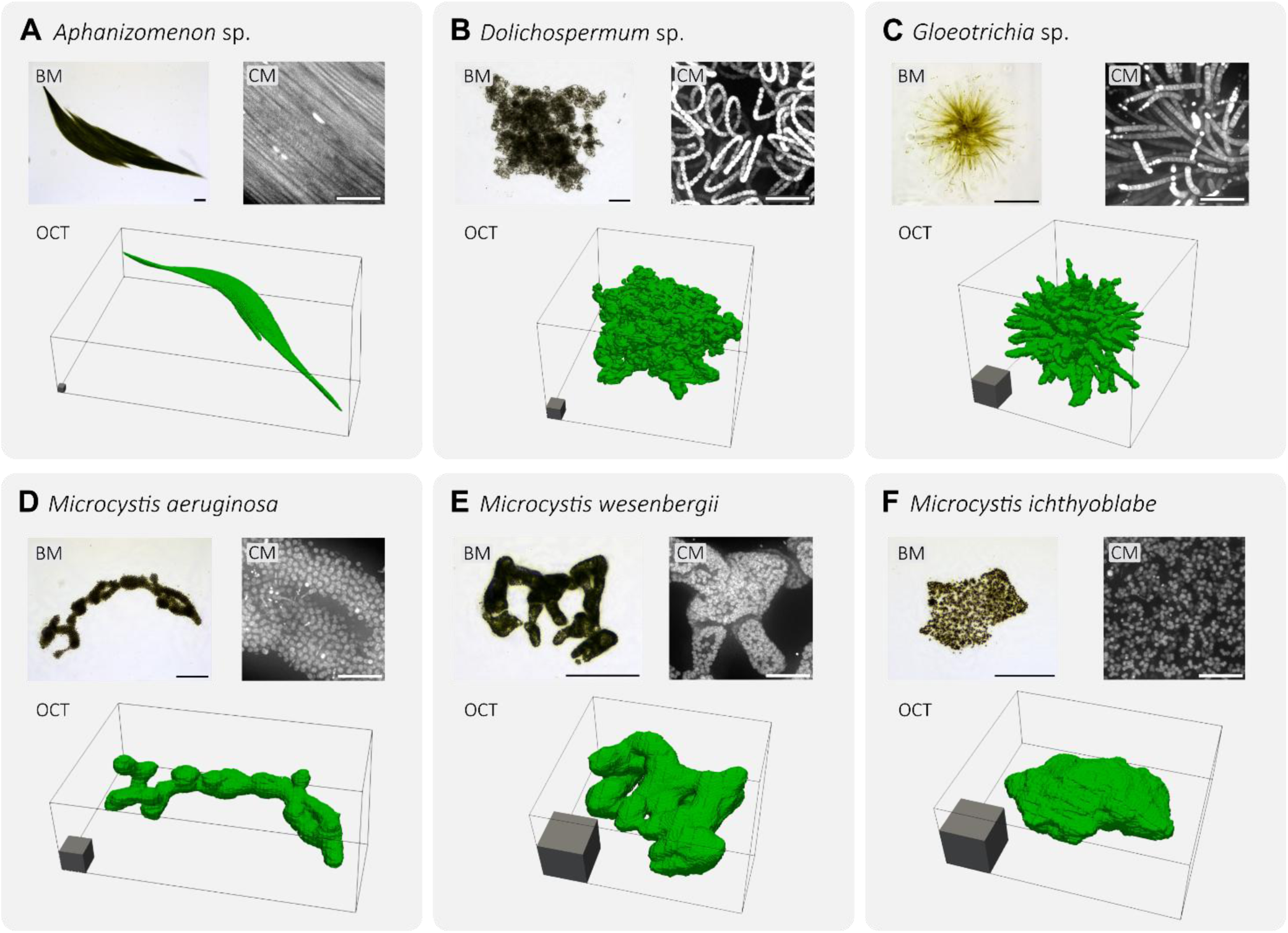
Volumetric visualization rendered from OCT scans of 6 colonies from different cyanobacterial species: (A) *Aphanizomenon* sp., (B) *Dolichospermum* sp., (C) *Gloeotrichia* sp., (D) *Microcystis aeruginosa*, (E) *M. wesenbergii*, (F) *M. ichthyoblabe*. Scale cube has 100 µm of side length. Top left insets for each panel display bright field microscopy (BM) images of the same colonies. Scale bars in the bright field images indicate 200 µm. Top right insets display maximum intensity projections of confocal fluorescence microscopy (CM) scans of the same colonies. Scale bars in the confocal images indicate 50 µm. Rotational views of the 3D structure of the colonies obtained by OCT are presented in Supplementary Movie S2.

Three different genera of cyanobacteria with filamentous colonies (Fig. 5A-C), namely *Aphanizomenon*, *Dolichospermum* and *Gloeotrichia*, and three morphospecies of *Microcystis* colonies (Fig. 5D-F) were imaged. A colony of *Aphanizomenon* sp. was formed of tightly arranged parallel filaments that formed a raft-like structure (Fig. 5A). Individual filaments were not distinguishable from the OCT scan, as the spacing between filaments was smaller than the lateral resolution of the OCT measurements. Thus, the colony appears in the volumetric visualization as a wide and thin continuous structure. During the experiments the filaments were actively gliding, therefore dynamically changing the shape of the colony, mainly in the lateral direction. The maximum displacement of filaments was smaller than the OCT lateral resolution (13 µm), thus the colony can be considered static with respect to the OCT acquisition time. A colony of *Dolichospermum* sp. was composed of coiled filaments entangled with each other (Fig. 5B). Similar to the colony of *Aphanizomenon* sp., individual filaments within the colony of *Dolichospermum* sp. were not distinguishable from the volumetric visualization. However, the spacing between filaments was in some regions larger than the OCT lateral resolution, thus the volumetric visualization of the colony was highly porous and corrugated. A colony of *Gloeotrichia* sp. was spherical and composed of straight filaments radiating from the colony center (Fig. 5C). The tips of each filament were distinguishable from the volumetric visualization, though their bases were merged due to the tight packing.

For all three morphospecies of *Microcystis* colonies, the individual cells were not distinguishable from the volumetric visualization (Fig. 5D-F). Moreover, the spacing between neighboring cells was smaller than the OCT lateral resolution. Therefore, the volumetric visualization of the *Microcystis* colonies displayed a continuous region corresponding to the cells and small intercellular voids. Microscopy visualization of EPS stained with Alcian Blue 8GX indicated that the intercellular voids in *Microcystis* colonies were filled with EPS (Fig. S3), in line with previous studies (Shen et al., 2011; Sinzato et al., 2026b). A branched structure was observed from the volumetric visualization of the morphospecies *Microcystis aeruginosa* (Fig. 5D) and *Microcystis wesenbergii* (Fig. 5E), while a flattened morphology was observed from the volumetric visualization of the morphospecies *Microcystis ichthyoblabe* (Fig. 5F). The refractive EPS matrix characteristic of *Microcystis wesenbergii* was only discernible from cells in the bright field microscopy image.

### C. Analysis of morphological features of *Microcystis* spp. colonies

Here we present mesoscale morphological features measured from the 3D binary images of *Microcystis* spp. colonies sampled from Lake ‘t Joppe in September 2024 (J-09-24) and August 2025 (J-08-25), and compare the results with an idealized spherical colony. In contrast to the common characterization of colonies with wide field microscopy, which essentially measures colony size from a 2D image projection, the 3D binary image acquired by OCT directly provides the colony volume. Our results indicate that the 2D image projection along the *z*-axis systematically overestimates the colony volume, as the equivalent circular diameter of each colony was larger than its equivalent spherical diameter (Fig. 6A). Both size descriptors were related by a power-law, *D_es_*⁄*L_d_* = (*D_ec_*⁄*L_d_*)*^f^_d_*, where *L_d_* = 61 ± 9 µm and *f_d_* = 0.83 ± 0.02 (mean ± SD, *R*^2^ = 0.92, *N* = 213 for the combined samples; dashed line in Fig. 6A), indicating that the deviation from a spherical shape (*D_es_* = *D_ec_* ; solid line in Fig. 6A) increases with colony size.

**Fig. 6:**
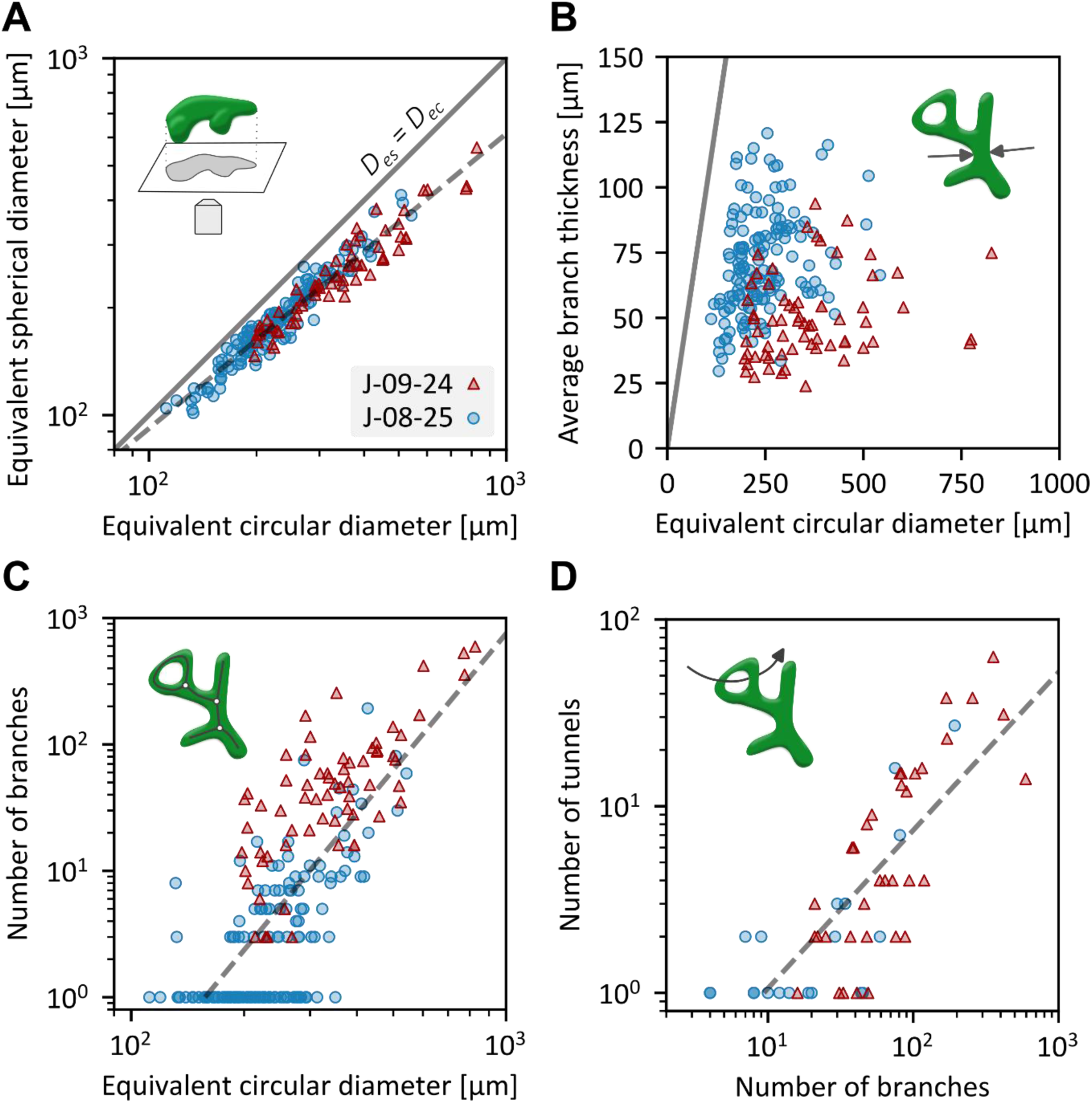
Quantitative morphological analysis of three-dimensional binary images of *Microcystis* spp. colonies acquired by OCT. (A) Relation between the equivalent spherical diameter *D_es_* obtained from the 3D colony volume and the equivalent circular diameter *D_ec_* measured from the 2D projection of the colony along the *z*-axis. (B) Average thickness of all branches within a colony as a function of the equivalent circular diameter. (C) Number of branches within each colony as a function of the equivalent circular diameter. (D) Number of tunnels within each colony as a function of the number of branches in colonies with at least one tunnel. Solid line in panels A and B indicates the predictions for an idealized spherical colony. Dashed lines in panels A, C and D indicate a power-law fit for the combined samples (see Supplementary Table S2 for fit parameters). Data points in all panels indicate individual colonies sampled from Lake ‘t Joppe in September 2024 (sample J-09-24 (triangles): *N* = 65 in panels A-C and *N* = 35 in panel D) and August 2025 (sample J-08-25 (bullets): *N* = 148 in panels A-C and *N* = 22 in panel D).

The overestimation of colony volume from its 2D projection indicated that large colonies had an irregular shape. We quantified colony shape using morphological skeleton analysis. An ideal spherical compact colony has a single point branch, with its thickness equal to the sphere diameter. However, the average branch thickness of the colonies was not significantly correlated with their equivalent circular diameter (Pearson correlation : *r*(211) = -0.01, *p =* 0.86, Fig. 6B), with a mean value (± SD) between all colonies of 65 ± 23 µm. Two main morphological routes can be identified for a colony to increase in size while maintaining a nearly constant branch thickness. Flattened compact colonies, typical of the morphospecies *M. ichthyoblabe* (Fig. 5F), have only one or a few branches and remain short in length in the axial direction, limiting their thickness while expanding their 2D projection. Another morphological route is that of branched colonies, typical of the morphospecies *M. aeruginosa* (Fig. 5D). A morphology with many branches allows the colony to increase its projected area while limiting the increase in volume and branch thickness. The number of branches measured per individual colony was positively correlated with its equivalent circular diameter (Pearson correlation using the log-transformed values: *r*(211) = 0.76, *p <* 0.001, Fig. 6C), with several hundred branches in large colonies, suggesting that colonies transition from compact to branched structures as they grow larger.

A highly branched structure also favored the formation of tunnels within colonies, as the number of tunnels was positively correlated with the number of branches (Pearson correlation for colonies with at least one tunnel, using the log-transformed values: *r*(55) = 0.80, *p <* 0.001, Fig. 6D). Void formation was more commonly observed in large colonies, as the number of voids was positively correlated with the equivalent circular diameter of the colonies (Pearson correlation for colonies with at least one void, using the log-transformed values: *r*(51) = 0.60, *p <* 0.001, Fig. S4). However, voids in all colonies were small, with a maximum ratio of void volume per colony volume of 2%, and thus had little impact on the outer shape of colonies.

Comparison between the 2024 and 2025 samples of Lake ‘t Joppe indicates that colonies from the 2024 sample had a larger average size, accompanied by a larger number of branches and tunnels and a smaller branch thickness (Fig. 6). This difference is likely due to the sampling time within the season and morphospecies composition of each sample. The 2025 sample was collected in early August 2025 and was dominated by *M. ichthyoblabe* colonies (59%), with *M. aeruginosa* (18%), *M. flos-aquae* (17%) and *M. wesenbergii* (6%) comprising the remainder. In contrast, the 2024 sample was collected in early September 2024 and contained similar proportions of *M. ichthyoblabe* (45%) and *M. aeruginosa* (43%), with the remaining fraction composed of *M. flos-aquae* (12%).

We next demonstrate how the measured 3D morphologies directly impact two key ecological traits: flotation velocity of the colonies and light availability within the colonies. We exemplified the effect of irregular colony morphologies by using the volumetric images of the dataset of *Microcystis* spp. colonies to model their flotation velocity and intracolony light attenuation. According to the classic Stokes’ law, the flotation velocity of an idealized spherical colony increases with the square of its diameter (Eq. 1 with a shape factor *ψ* = 1). Instead, estimates of the shape factor of the scanned colonies (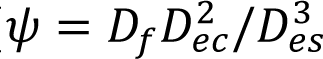; Sinzato et al., 2026b) increased systematically with their equivalent circular diameter following a power law (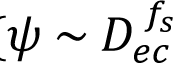, Fig. S5) with an exponent of *f_s_* = 0.67 ± 0.06 (mean ± SD, combined samples, Supplementary Table S2). A comparison between the individual samples of Lake ‘t Joppe indicates that the power law for the 2024 sample had a higher exponent (*f_s_* = 0.75 ± 0.13) compared to 2025 sample (*f_s_* = 0.43 ± 0.07), which likely reflects the larger number of branches and tunnels in the 2024 sample. The power law for the 2024 sample obtained here from the 3D scans of colonies agrees well, within the uncertainty range, with that obtained from colony tracking experiments in our previous work, which used samples collected at the same location and time (*f_s_* = 0.82 ± 0.04; Sinzato et al., 2026b). Implementation of the size-dependence of the shape factor in Eq. (1) results in a scaling of the flotation velocity with 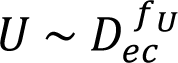, where *f_U_* = 2 − *f_s_* = 1.33 ± 0.06 for the combined samples (dashed line in Fig. 7A). That is, the flotation velocity of *Microcystis* colonies does not scale with the square of colony diameter as in classic Stokes’ law, but with a smaller exponent of 1.33. This results in a smaller flotation velocity compared to that of a spherical colony (solid line in Fig. 7A).

**Fig. 7:**
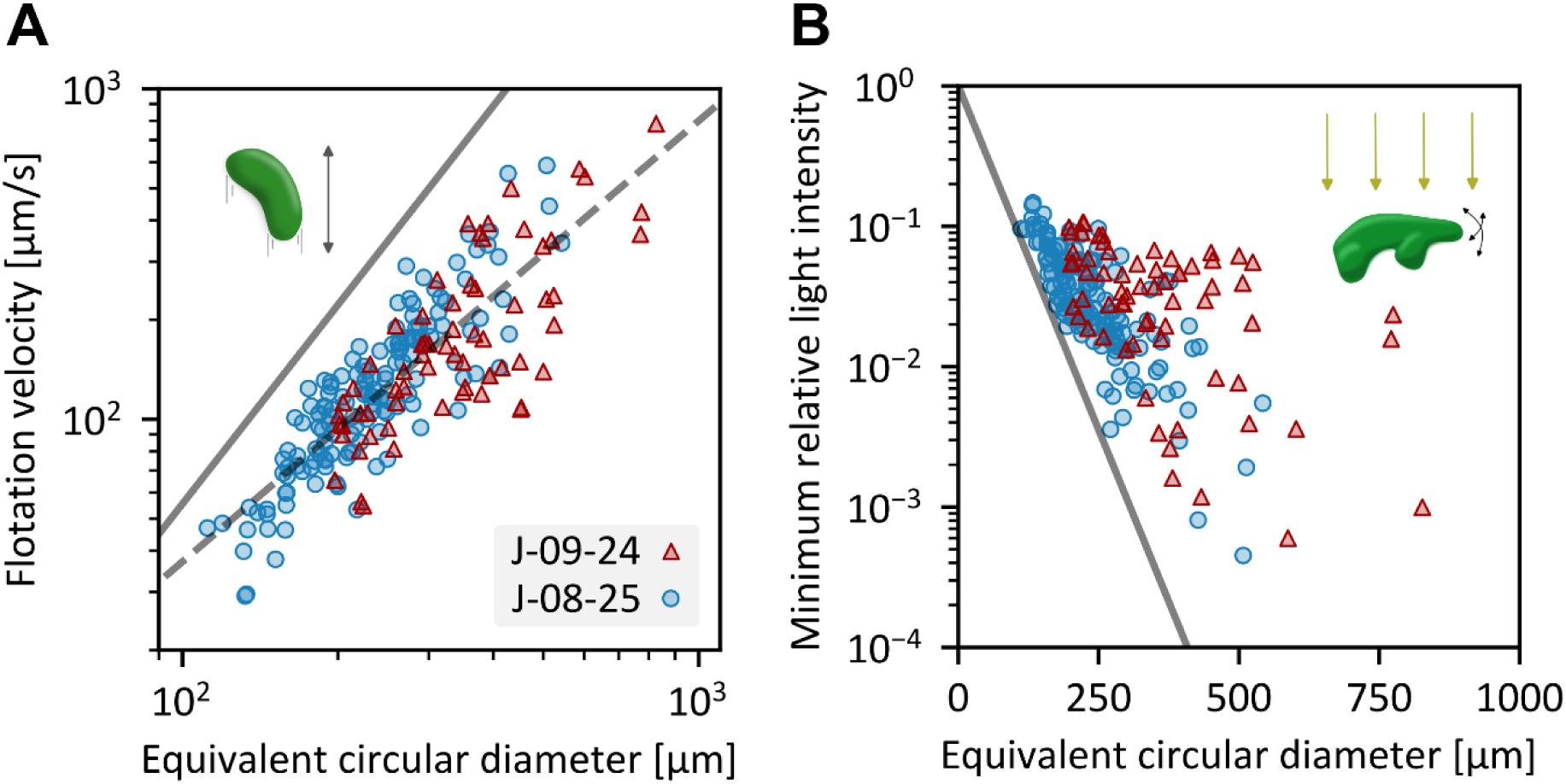
Effect of three-dimensional colony morphologies on two key ecological traits. (A) Estimated flotation velocity as a function of the equivalent circular diameter. (B) Minimum value of the relative light intensity inside a colony estimated from the Lambert-Beer law and averaged over all orientations. Solid lines in each panel indicate the predictions for an idealized spherical colony. Dashed line in panel A indicates Eq. (1) with the power law fit for the shape factor (see Supplementary Table S2 for fit parameters). Data points in all panels indicate individual colonies sampled from Lake ‘t Joppe in September 2024 (sample J-09-24 (triangles): *N* = 65) and August 2025 (sample J-08-25 (bullets): *N* = 148).

Light availability within the colony was modeled according to Eq. (4). For each colony, the minimum relative light intensity (normalized by the incident light intensity) within the colony was computed (Fig. 7B). Strong light attenuation was present in the center of large colonies, reaching minimum relative light intensities of *I_min_/I_o_ <* 0.01 for some colonies. However, the irregular shapes of the colonies allowed them to achieve minimum relative light intensities much higher than the theoretical value for spherical colonies (*I_min_/I_o_* = exp(−*ε D_ec_ /* 2), solid line in Fig. 7B). Similar conclusions were also obtained for the relative light intensity averaged over the entire volume of each colony (Fig. S6). This result indicates that, compared to spherical colonies, a branched colony structure enhances light availability within the colony.

## IV. DISCUSSION

Our results show that OCT image acquisition and an automated image processing pipeline can be successfully employed to obtain 3D visualizations of colonies of different cyanobacterial species. The technique offers several advantages over existing imaging methods. The imaging depth and field of view were sufficient to fully scan the entire colonies (maximum colony dimensions were 9 mm in length and 1 mm in height), which is a major advantage over confocal fluorescence microscopy. Moreover, the sample preparation steps are minimal, can be applied on both fresh and fixed samples, and are comparatively easier than those required for transmission electron microscopy. The fast acquisition time is another advantage of OCT, as it allows for a high acquisition throughput and the imaging of cyanobacteria with motile filaments such as *Aphanizomenon*. The presence of gas vesicles inside cells strongly enhanced the signal intensity of colonies, although collapsed gas vesicles still allowed for sufficient contrast between the colony and background.

Mesoscale morphological features of ∼200 *Microcystis* spp. colonies were obtained. The results from this extensive dataset can be used in other studies. For example, the power-law relation between the equivalent spherical diameter and the equivalent circular diameter of the colonies provides a correction for projection-bias in estimates of the biovolume of cyanobacterial blooms. The skeleton analysis presented here offers a method to quantify differences in shape between different morphospecies of *Microcystis*, and can potentially be implemented in future studies on colony formation and seasonal succession in *Microcystis* populations. The 3D images of the colonies in combination with a modified Stokes’ law provide estimates of the shape factor used to calculate their flotation velocity, without the need for controlled velocity measurements of colonies tracked in laboratory experiments. The estimated shape factor can be readily implemented in models predicting the vertical distribution of cyanobacteria during the formation of harmful blooms (Huisman et al., 2004; Ranjbar et al., 2024; Feng et al., 2025). Finally, a model of light attenuation within the colony revealed that the branched shapes of large colonies can strongly improve light availability in inner cells compared to spherical colonies. This result corroborates previous experimental findings that large *Microcystis* colonies can adjust their cell arrangement to avoid excessive self-shading (Feng et al., 2024). Although our model neglects light scattering and variations in cell concentration within colonies, such effects could be incorporated in future studies to understand colony growth under light-limited conditions (Feng et al., 2024).

An interesting observation from our OCT analysis is that the deviation from a spherical shape (Fig. 6A) as well as the number of branches, voids and tunnels (Fig. 6C,D; Fig. S4) all increased with colony size. Since colony formation in natural *Microcystis* populations is largely dominated by cell division (Pérez-Carrascal et al., 2021; Smith et al., 2021; Sinzato et al. 2026a), these results indicate that the morphology of *Microcystis* colonies, which is commonly used for the classification of different morphospecies, alters from compact near-spherical shapes towards more branched nonspherical morphotypes as colonies grow larger during the summer season. This aligns with the increase in colony size and the shift in morphotype composition of the *Microcystis* colonies we collected from Lake ‘t Joppe in early August 2025 and early September 2024. Hence, our results provide support for a transition route from *M. ichthyoblabe*-like to *M. aeruginosa-*like colonies, with *M. wesenbergii* as an intermediate morphology, previously proposed to explain seasonal variation in colony morphology (Li et al., 2013; Xiao et al., 2018).

A major limitation of OCT measurements is the poor lateral resolution, which is insufficient to resolve single cells and filaments within cyanobacterial colonies. Consequently, the colony volume measured with OCT also incorporates intercellular or inter-filament regions. These regions may be water-filled or EPS-filled, and the conversion of colony volume to cell number within a colony still requires additional optical microscopy. This limitation in the resolution of classic OCT can potentially be improved with the development of micro-optical coherence tomography (Nishimiya & Tearney, 2021). While the experiments presented here were conducted on freshwater cyanobacteria, the methodology could be readily applied to marine cyanobacterial colonies, such as *Trichodesmium* (Capone et al., 1997). We expect that optical coherence tomography will greatly advance future research on colony formation in cyanobacteria as well as in other colonial plankton taxa and suspended biological aggregates.

## V. COMMENTS AND RECOMMENDATIONS

The sample preparation protocol presented here can be implemented with other imaging chambers, such as a Sedgewick-Rafter chamber, 96-well microplate, or custom-made geometries. Attention should be given to the reflection of the chamber surfaces: strong reflections introduce significant noise, yet a small amount of reflection is required to detect the surfaces of the imaging chamber and correct for chamber tilt. The tilt angle (either of the chamber or of the OCT scanner) can be adjusted to find an optimal configuration. The operator should also take care to avoid bubbles in the chamber and to centre the sample within the field of view.

Although it is preferable to keep gas vesicles intact for OCT imaging, some preparation procedures such as EPS staining require their collapse. In such cases, the signal-to-noise ratio can be improved by averaging several scans or by increasing scanner sensitivity, at the cost of longer acquisition times. Conversely, imaging of highly motile colonies or colonies in a flow chamber may require faster acquisition, accepting a weaker signal as a trade-off.

## ACKNOWLEDGMENTS

We are grateful to Axel Gunderson for his assistance with fieldwork. We thank Nico Schramma and Isaac Deen Garcia for their support with image processing. We especially thank Johan Oosterbaan for his contributions during the project’s conception. We acknowledge funding from the Hoogheemraadschap van Rijnland (Rijnland Regional Water Authority). MJ acknowledges support from the ERC grant no. 2023-StG-101117025, FluMAB.

## DATA AND CODE AVAILABILITY

Primary data of OCT scans, and the dataset of binary images and measurements of mesoscale features for *Microcystis* spp. colonies are available at https://doi.org/10.21942/uva.33329520. The script for image processing is available at https://github.com/FluidLab/CyanOCT.

## SUPPORTING INFORMATION FOR

### Supplementary Tables

**Table S1:** Location and date of collection for the reported images of cyanobacterial colonies.

| Figure | Species | Location | Date |
| --- | --- | --- | --- |
| Fig. 1 | *Microcystis aeruginosa* | Lake Haarlemmermeerse Bosplas | 6 Oct 2025 |
| Fig. 2 and 3 | *Microcystis ichthyoblabe* | Lake Braassemermeer | 18 Jul 2025 |
| Fig. 4 | *Dolichospermum* sp. | Lake Haarlemmermeerse Bosplas | 18 Jun 2025 |
| Fig. 5A | *Aphanizomenon* sp. | Lake 't Joppe | 7 Aug 2025 |
| Fig. 5B | *Dolichospermum* sp. | Lake Haarlemmermeerse Bosplas | 18 Jun 2025 |
| Fig. 5C | *Gloeotrichia* sp. | Lake Gaasperplas | 10 Jun 2025 |
| Fig. 5D | *Microcystis aeruginosa* | Lake 't Joppe | 7 Aug 2025 |
| Fig. 5E | *Microcystis wesenbergii* | Lake 't Joppe | 7 Aug 2025 |
| Fig. 5F | *Microcystis ichthyoblabe* | Lake Braassemermeer | 18 Jul 2025 |
| Fig. 6 and 7 | *Microcystis* spp. (sample J-08-25) | Lake 't Joppe | 7 Aug 2025 |
| Fig. 6 and 7 | *Microcystis* spp. (sample J-09-24) | Lake 't Joppe | 3 Sep 2024 |
| Fig. S1 | *Microcystis aeruginosa* | Lake 't Joppe | 7 Aug 2025 |
| Fig. S2 | *Microcystis ichthyoblabe* | Lake Braassemermeer | 18 Jul 2025 |
| Fig. S3 | *Microcystis ichthyoblabe* | Lake Braassemermeer | 18 Jul 2025 |
| Fig. S4, S5, S6 | *Microcystis* spp. (sample J-08-25) | Lake 't Joppe | 7 Aug 2025 |
| Fig. S4, S5, S6 | *Microcystis* spp. (sample J-09-24) | Lake 't Joppe | 3 Sep 2024 |

**Table S2:** Correlation coefficients and fit parameters between morphological features for the individual and combined samples. Fit parameters are reported as mean ± SD. Power-law fits are based on the log-transformed data values.

| Feature Y | Feature X | Pearson correlation | Power law fit |
| --- | --- | --- | --- |
| Equivalent spherical diameter $D_{es}$ | Equivalent circular diameter $D_{ec}$ | J-09-24: $r^*(63)=0.95, p<0.001$<br>J-08-25: $r^*(143)=0.97, p<0.001$<br>All: $r^*(211)=0.97, p<0.001$ | $D_{es} = L_d(D_{ec}/L_d)^{f_d}$<br>J-09-24: $L_d = 75 \pm 18 \mu\text{m}; f_d = 0.79 \pm 0.03$<br>J-08-25: $L_d = 33 \pm 11 \mu\text{m}; f_d = 0.89 \pm 0.02$<br>All: $L_d = 61 \pm 9 \mu\text{m}; f_d = 0.83 \pm 0.02$ |
| Average branch thickness $H_b$ | Equivalent circular diameter $D_{ec}$ | J-09-24: $r(63)=0.23, p=0.07$<br>J-08-25: $r(143)=0.29, p=0.003$<br>All: $r(211)=-0.01, p=0.86$ | No significant correlation. Mean value:<br>J-09-24: $\langle H_b \rangle = 50 \pm 16 \mu\text{m}$<br>J-08-25: $\langle H_b \rangle = 72 \pm 22 \mu\text{m}$<br>All: $\langle H_b \rangle = 65 \pm 23 \mu\text{m}$ |
| Number of branches $N_b$ | Equivalent circular diameter $D_{ec}$ | J-09-24: $r^*(63)=0.72, p<0.001$<br>J-08-25: $r^*(143)=0.67, p<0.001$<br>All: $r^*(211)=0.76, p<0.001$ | $N_b = (D_{ec}/L_d)^{f_d}$<br>J-09-24: $L_d = 76 \pm 14 \mu\text{m}; f_d = 2.4 \pm 0.3$<br>J-08-25: $L_d = 160 \pm 7 \mu\text{m}; f_d = 2.6 \pm 0.2$<br>All: $L_d = 157 \pm 6 \mu\text{m}; f_d = 3.6 \pm 0.2$ |
| Number of tunnels $N_t$ | Number of branches $N_b$ | J-09-24: $r^*(33)=0.78, p<0.001$<br>J-08-25: $r^*(20)=0.73, p<0.001$<br>All: $r^*(55)=0.80, p<0.001$ | $N_t = (N_b/N_o)^{f_d}$<br>J-09-24: $N_o = 15 \pm 4; f_d = 1.07 \pm 0.15$<br>J-08-25: $N_o = 7 \pm 2; f_d = 0.64 \pm 0.13$<br>All: $N_o = 9 \pm 2; f_d = 0.85 \pm 0.09$ |
| Number of voids $N_v$ | Equivalent circular diameter $D_{ec}$ | J-09-24: $r^*(33)=0.58, p<0.001$<br>J-08-25: $r^*(20)=0.62, p=0.002$<br>All: $r^*(55)=0.59, p<0.001$ | $N_v = (D_{ec}/L_d)^{f_d}$<br>J-09-24: $L_d = 200 \pm 36 \mu\text{m}; f_d = 1.7 \pm 0.4$<br>J-08-25: $L_d = 237 \pm 29 \mu\text{m}; f_d = 2.2 \pm 0.6$<br>All: $L_d = 213 \pm 24 \mu\text{m}; f_d = 1.8 \pm 0.3$ |
| Shape factor $\psi$ | Equivalent circular diameter $D_{ec}$ | J-09-24: $r^*(63)=0.57, p<0.001$<br>J-08-25: $r^*(143)=0.51, p<0.001$<br>All: $r^*(211)=0.61, p<0.001$ | $\psi = (D_{ec}/L_s)^{f_s}$<br>J-09-24: $L_s = 59 \pm 19 \mu\text{m}; f_s = 0.75 \pm 0.13$<br>J-08-25: $L_s = 28 \pm 10 \mu\text{m}; f_s = 0.43 \pm 0.07$<br>All: $L_s = 55 \pm 8 \mu\text{m}; f_s = 0.67 \pm 0.06$ |

\* Correlation between the log-transformed values

### Supplementary Figures

**Fig. S1:**
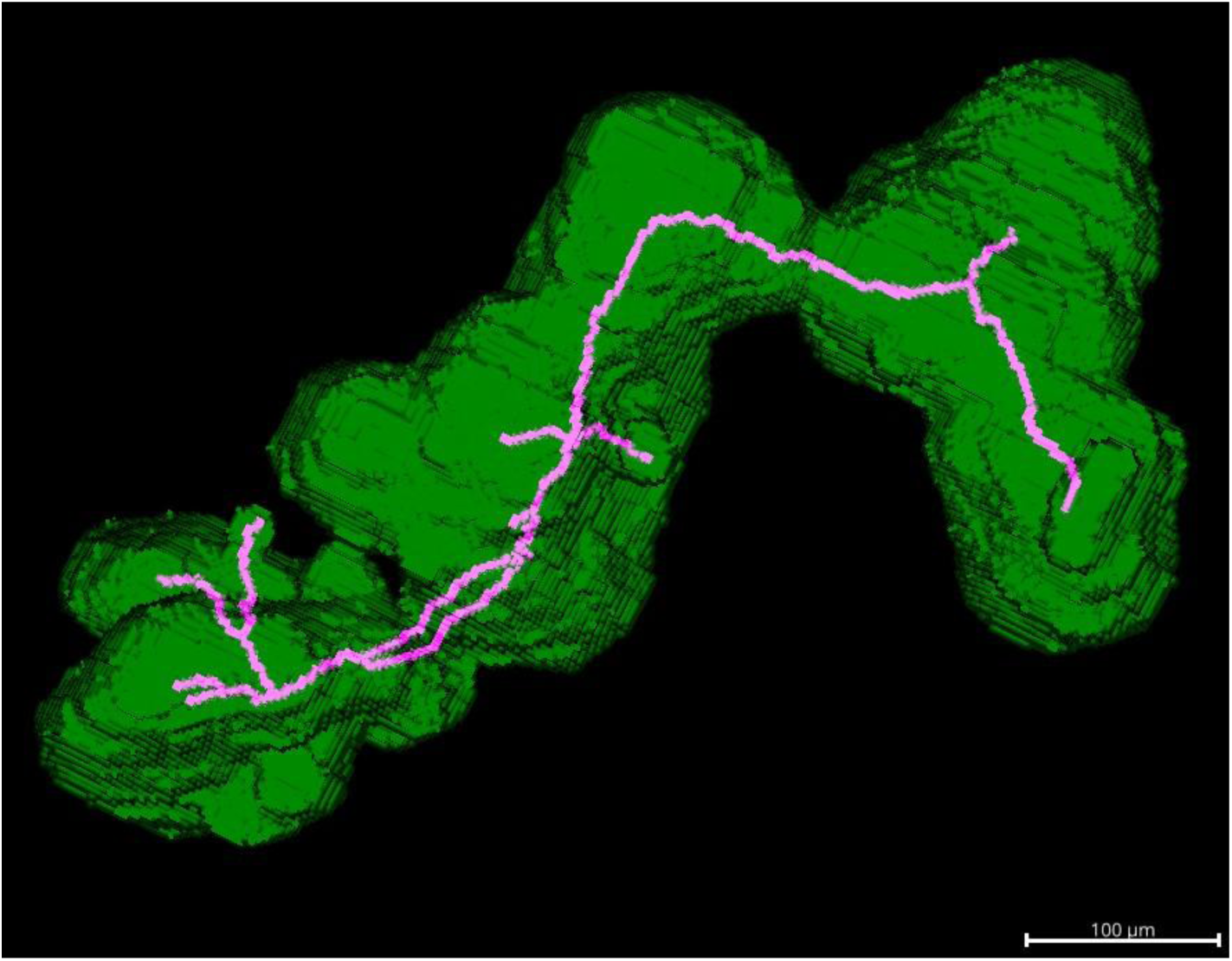
Three-dimensional visualization of the skeleton (magenta line) of a *Microcystis* colony overlapped with its volumetric visualization (green surface). A rotational view of the 3D structure is presented in Supplementary Movie S1.

**Fig. S2:**
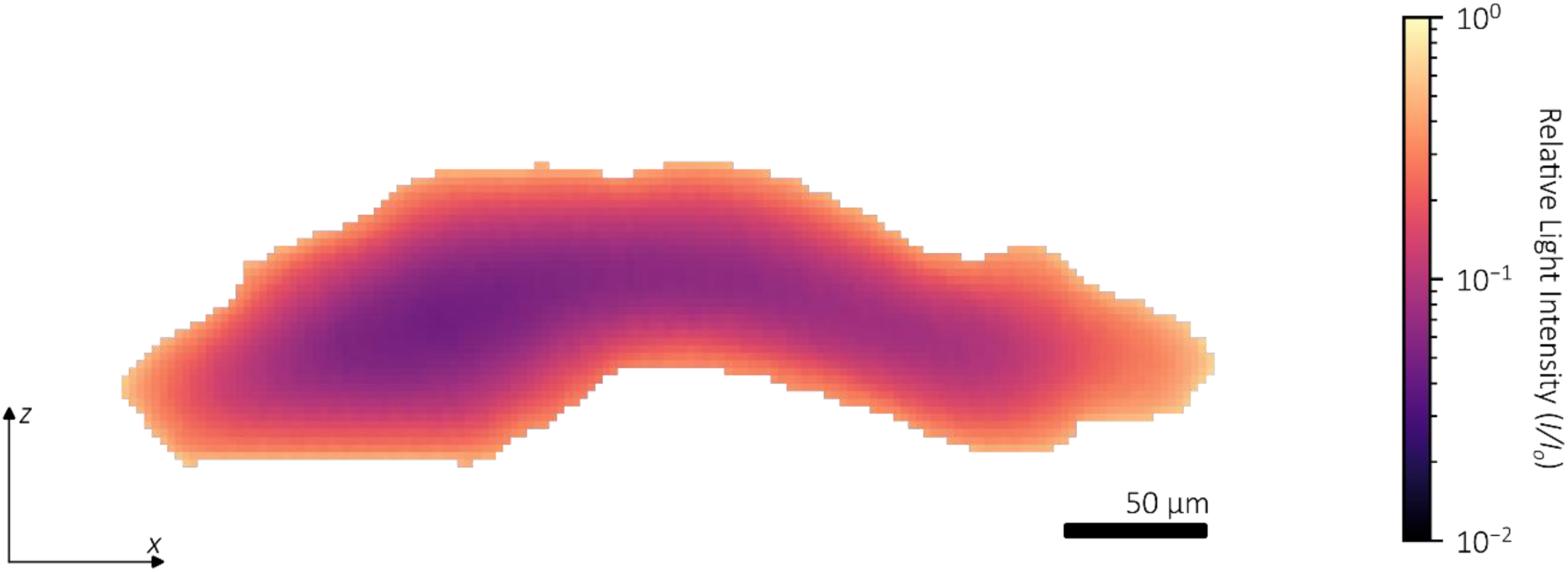
Relative light intensity inside a slice of a *Microcystis* colony following light attenuation computed from Eq. (4).

**Fig. S3:**
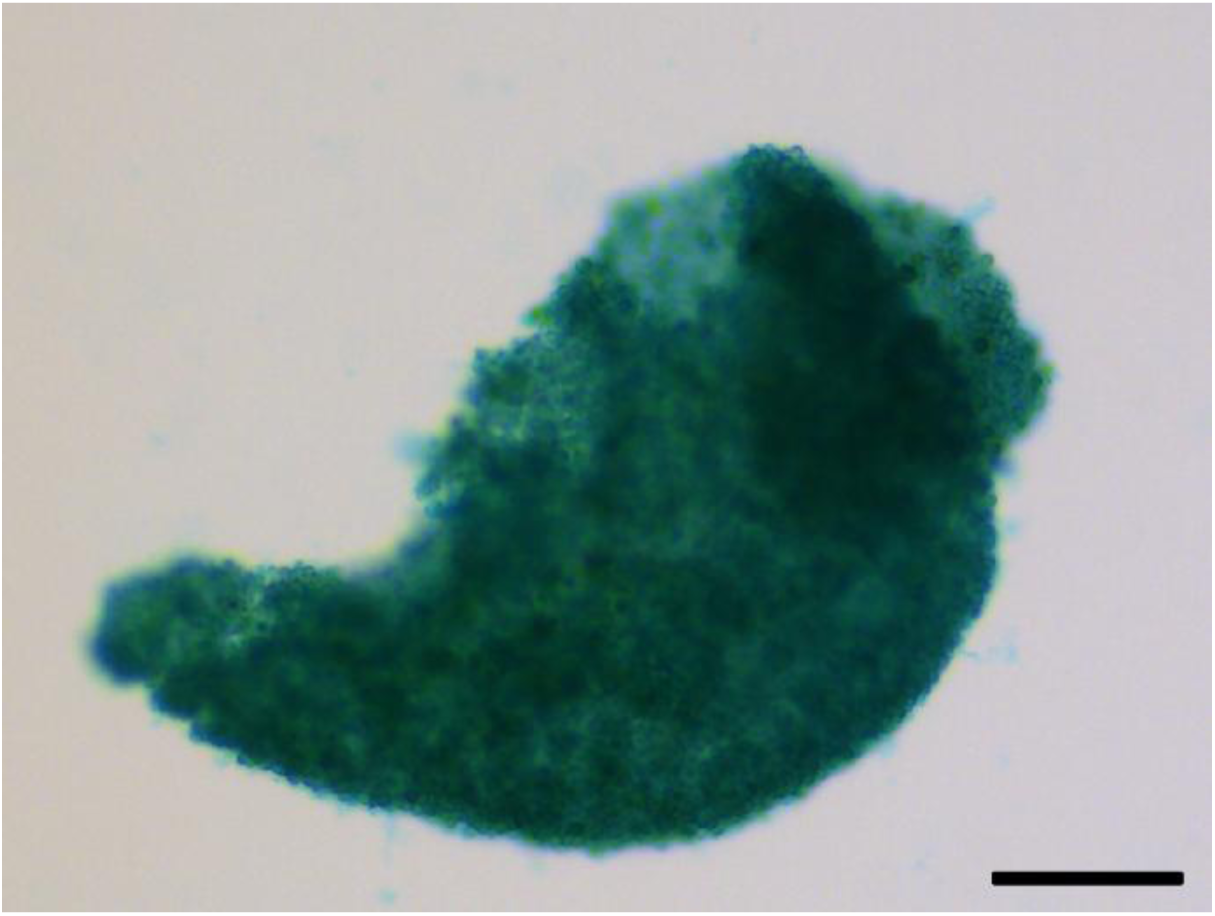
Bright field microscopy image of a *Microcystis* colony with EPS stained with Alcian Blue 8GX. The colony with collapsed gas vesicles is incubated for 30 min in 0.3 % (w/v) Alcian Blue 8GX in 3 % (v/v) acetic acid. After incubation, the excess staining solution is removed, the colony is dispersed in 3 % (v/v) acetic acid, then pipetted into an imaging chamber. Stained EPS appears blue. Scale bar indicates 100 µm.

**Fig. S4:**
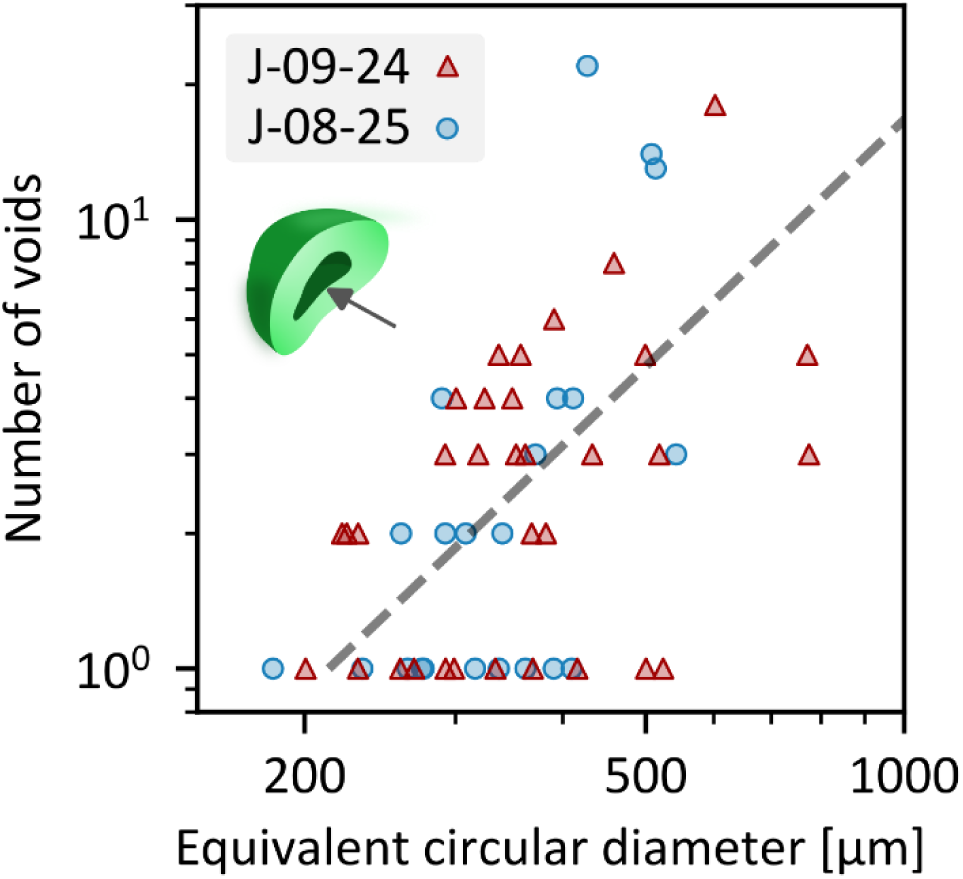
Number of voids inside *Microcystis* spp. colonies as a function of the equivalent circular diameter. Pearson correlation using the log-transformed number of voids and equivalent circular diameter of colonies with at least one void: *r*(55) = 0.60, *p* < 0.001. Dashed line indicates a power-law fit (see Supplementary Table S2 for fit parameters). Data points in all panels indicate individual colonies sampled from Lake ‘t Joppe in September 2024 (sample J-09-24 (triangles): *N* = 35) and August 2025 (sample J-08-25 (bullets): *N* = 22).

**Fig. S5:**
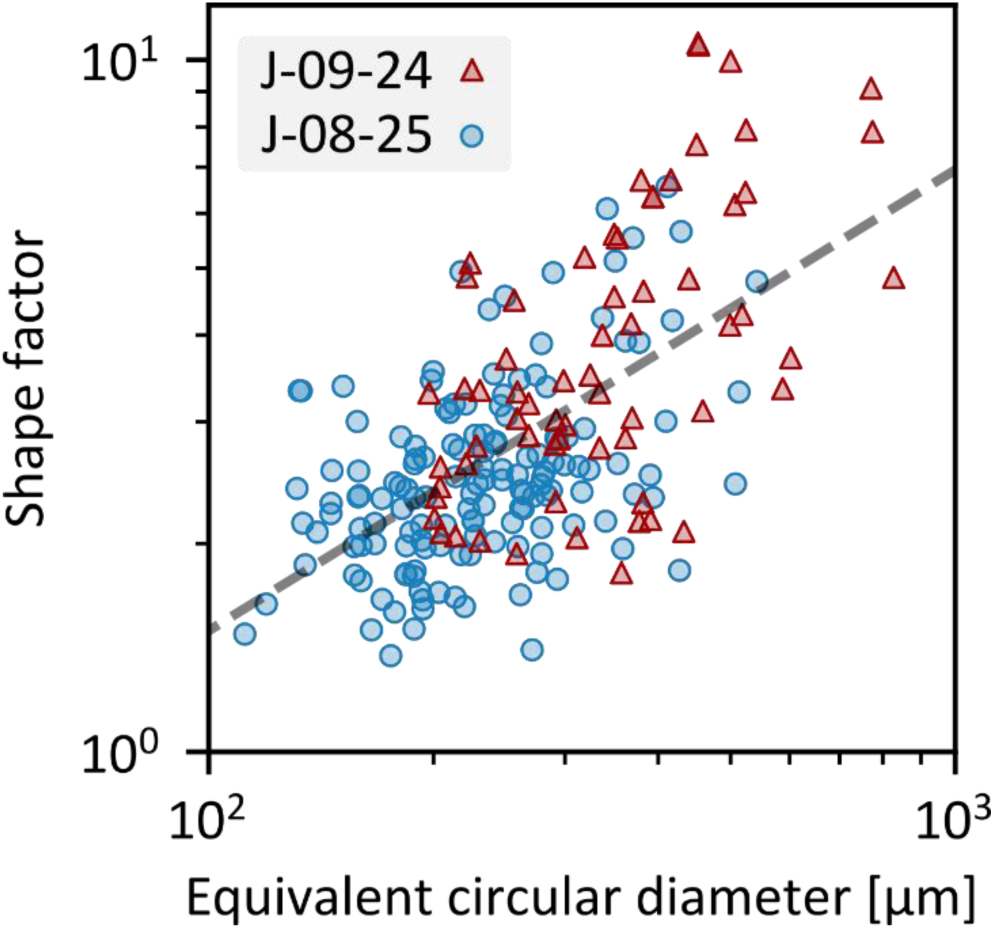
Shape factor of *Microcystis* spp. colonies as a function of the equivalent circular diameter. Pearson correlation using the log-transformed shape factor and equivalent circular diameter of colonies: *r*(211) = 0.61, *p* < 0.001. Dashed line indicates a power-law fit (see Supplementary Table S2 for fit parameters). Data points indicate individual colonies sampled from Lake ‘t Joppe in September 2024 (sample J-09-24 (triangles): N = 65) and August 2025 (sample J-08-25 (bullets): N = 148).

**Fig. S6:**
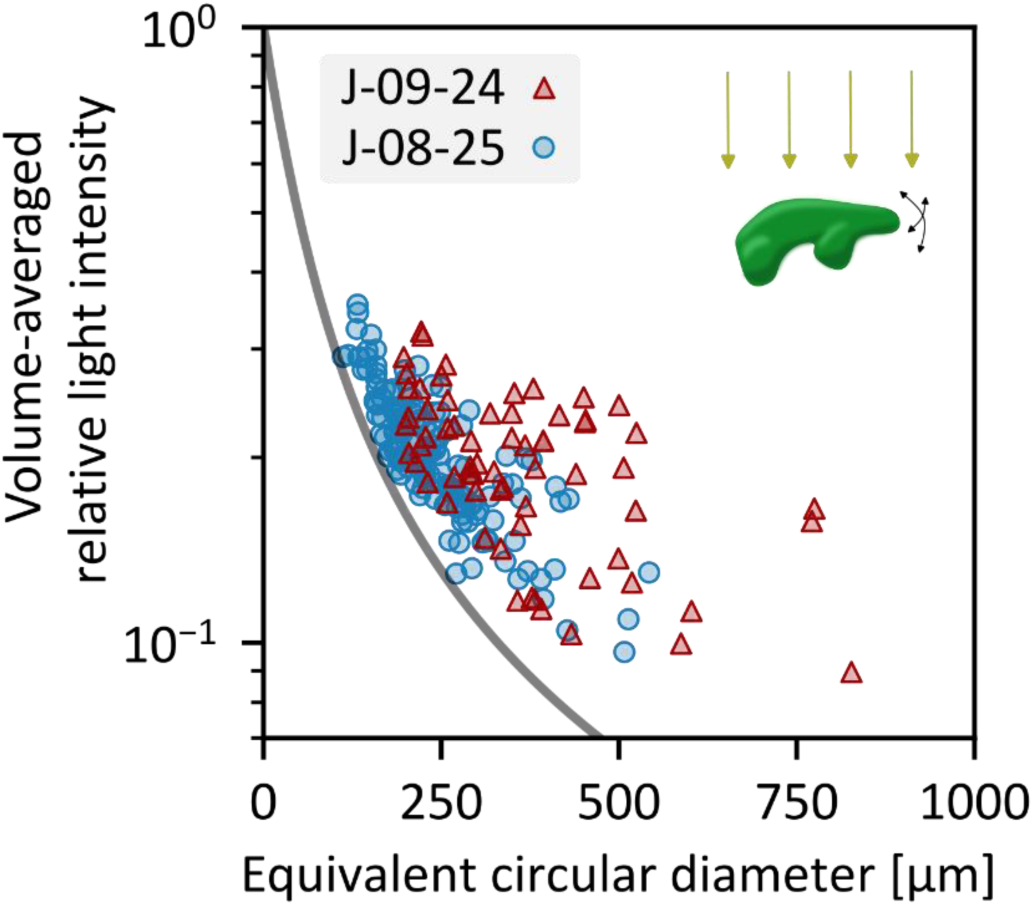
Volume-averaged relative light intensity: Light intensity inside a colony was estimated from Lambert-Beer law and averaged over all orientations and over the entire colony volume. Solid line indicates the prediction for an idealized spherical colony: 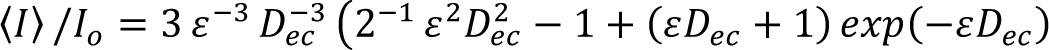. Data points indicate individual colonies sampled from Lake ‘t Joppe in September 2024 (sample J-09-24 (triangles): N = 65) and August 2025 (sample J-08-25 (bullets): N = 148).

Supplementary movies are available in the online version:

- Supplementary Movie S1 – Rotational view of the 3D structure of the skeleton and volumetric visualization of a *Microcystis* colony obtained by OCT - Scale cube side length = 50 µm.
- Supplementary Movie S2 - Rotational views of the 3D structure of colonies of different cyanobacterial species obtained by OCT - Scale cube side length = 100 µm.

